# Machine-learning based detection of adventitious microbes in T-cell therapy cultures using long read sequencing

**DOI:** 10.1101/2022.11.03.514634

**Authors:** James P. B. Strutt, Meenubharathi Natarajan, Elizabeth Lee, Denise Bei Lin Teo, Wei-Xiang Sin, Faye Ka-Wai Cheung, Marvin Chew, Khaing Thazin, Paul W. Barone, Jacqueline M. Wolfrum, Rohan B. H. Williams, Scott A. Rice, Stacy L. Springs

**Affiliations:** Singapore-MIT Alliance for Research and Technology, Singapore; MIT Center for Biomedical Innovation, Massachusetts Institute of Technology, U.S.A.; CSIRO Microbiomes for One Systems Health, Agriculture and Food, Westmead, Australia; Singapore Centre for Environmental Life Sciences Engineering, Life Sciences Institute, National University of Singapore, Singapore; Singapore Centre for Environmental Life Sciences Engineering, Nanyang Technological University, Singapore

**Keywords:** T-cells, adventitious agents, machine learning, sterility

## Abstract

Assuring that cell therapy products are safe before releasing them for use in patients is critical. Currently, compendial sterility testing for bacteria and fungi can take 7-14 days. The goal of this work was to develop a rapid untargeted approach for the sensitive detection of microbial contaminants at low abundance from low volume samples during the manufacturing process of cell therapies. We developed a long-read sequencing methodology using Oxford Nanopore Technologies MinION platform with 16S and 18S amplicon sequencing to detect USP<71> organisms and other microbial species. Reads are classified metagenomically to predict the microbial species. We used an extreme gradient boosting machine learning algorithm (XGBoost) to first assess if a sample is contaminated and second, determine whether the predicted contaminant is correctly classified or misclassified. The model was used to make a final decision on the sterility status of the input sample. An optimised experimental and bioinformatics pipeline starting from spiked species through to sequenced reads allowed for the detection of microbial samples at 10 CFU / mL using metagenomic classification. Machine learning can be coupled with long read sequencing to detect and identify sample sterility status and microbial species present in T-cell cultures, including the USP<71> organisms to 10 CFU / mL.

**Importance:** This research presents a novel method for rapidly and accurately detecting microbial contaminants in cell therapy products, which is essential for ensuring patient safety. Traditional testing methods are time-consuming, taking 7-14 days, while our approach can significantly reduce this time. By combining advanced long read Nanopore sequencing techniques and machine learning, we can effectively identify the presence and types of microbial contaminants at low abundance levels. This breakthrough has the potential to improve the safety and efficiency of cell therapy manufacturing, leading to better patient outcomes and a more streamlined production process.

## Introduction

Cell therapies are increasingly prevalent in the treatment of incurable diseases. For example, chimeric antigen receptor T-cells (CAR-T) are used for the treatment of hematologic malignancies.^1^ On-going work with human pluripotent stem cells (hPSCs) is targeted to treat Parkinson’s and age-related macular degeneration (AMD),^2^ while mesenchymal stromal cells (MSCs), are being developed for immunomodulatory treatments.^3^ Compendial sterility methods based on microbial growth are laborious and slow, and faster methods are required to guide clinical management.^4^ Rapid testing methodologies could be an important tool for decreasing the time that a patient must wait from initial leukapheresis to application of the cell therapy. Depending on a patient’s current health status, the patient may not be able to afford delays in the application of a potentially lifesaving therapy. Thus, a reduction of release testing time will ensure the timely and safe delivery of life saving cell therapies leading to improved patient outcomes.

Current good manufacturing practice for microbial safety has been developed from experience in recombinant protein manufacturing, where standard practises includes three pillars of safety: 1) identifying appropriate sterile raw materials, 2) testing of cell banks and in-process microbe testing for materials, and 3) inclusion of process steps to inactivate and remove undetected microbes.^5^ Cell therapy manufacturers are currently only able to use the first two pillars, as the product cannot be terminally sterilised. For testing, As such, the main approach to microbial safety is with compendial sterility tests. For example, test samples are inoculated into multiple growth media that support proliferation of aerobic and anaerobic organisms, as well as using the plate-count method and membrane filtration.^6^ Validation of compendial methods includes testing with USP<71> organisms. For a list of USP<71> organisms used in this study, see Table 1. The procedures for ensuring that the tests give valid results are codified in the United States Pharmacopoeia / European Pharmacopoeia (USP/EP). These methodologies can detect contamination events but cannot determine the species identity, which requires additional time-consuming follow-up by the manufacturer for failure investigation. The USP<71> tests suffer from false negatives at low (< 30 CFU) contaminant concentration when organisms fail to grow to the point of visible detection in the allocated time.^4^ Alternative sterility testing approaches have been developed with lower limits of detection, such as, the BacT/ALERT system, which is an automated system that works by colorimetric change of CO_2_ level evolution monitored every 15 minutes. The BacT/ALERT system has a limit of detection of < 10 CFU for sample volumes of 0.5-10 mL.^7, 8^

**Table 1:** Culture conditions MRCB: Modified Reinforced Clostridial Broth; YPD: Yeast Extract–Peptone–Dextrose; BHIB: Brain Heart Infusion Broth; LB: Luria-Bertani broth; RT: room temperature; O/N: overnight. * Aspergillus brasilensis was purchased as a pellet from ATCC.

| Organism | Medium | Temperature | Culture time | Sub-culture time | Respiration |
| --- | --- | --- | --- | --- | --- |
| *Cutibacterium acnes* (ATCC-6919) | BHIB | 33°C | 5 d | 2 d | Anaerobic |
| *Klebsiella pneumoniae* (KP1) | LB | 37°C | O/N | 2 h | Aerobic |
| *Escherichia coli* (K12) | LB | 37°C | O/N | 2 h | Aerobic |
| *Pseudomonas aeruginosa* PAO1 (ATCC BAA47) | LB | 37°C | O/N | 2 h | Aerobic |
| *Candida albicans* (ATCC 10231) | YPD | RT | 2 d | 2 h | Aerobic |
| USP<71> organisms |  |  |  |  |  |
| *Staphylococcus aureus* subsp. Aureus (ATCC 6538) | LB | 37°C | O/N | 2 h | Aerobic |
| *Pseudomonas aeruginosa* (ATCC 9027) | LB | 37°C | O/N | 2 h | Aerobic |
| *Bacillus subtilis* subsp. Spizizenii (ATCC 6633) | LB/<br>BHIB | 33°C | O/N | 2 h | Aerobic |
| *Clostridium sporogenes* (ATCC 19404) | MRCB | 37°C | 2 d | 2 h | Anaerobic |
| *Candida albicans* (ATCC 10231) | YPD | RT | 2 d | 2 h | Aerobic |
| *Aspergillus brasiliensis* (ATCC 16404)* | - | - | - | - | Aerobic |

MRCB: Modified Reinforced Clostridial Broth; YPD: Yeast Extract–Peptone–Dextrose; BHIB: Brain Heart Infusion Broth; LB: Luria-Bertani broth; RT: room temperature; O/N: overnight. \* *Aspergillus brasiliensis* was purchased as a pellet from ATCC.

**Table 2:**
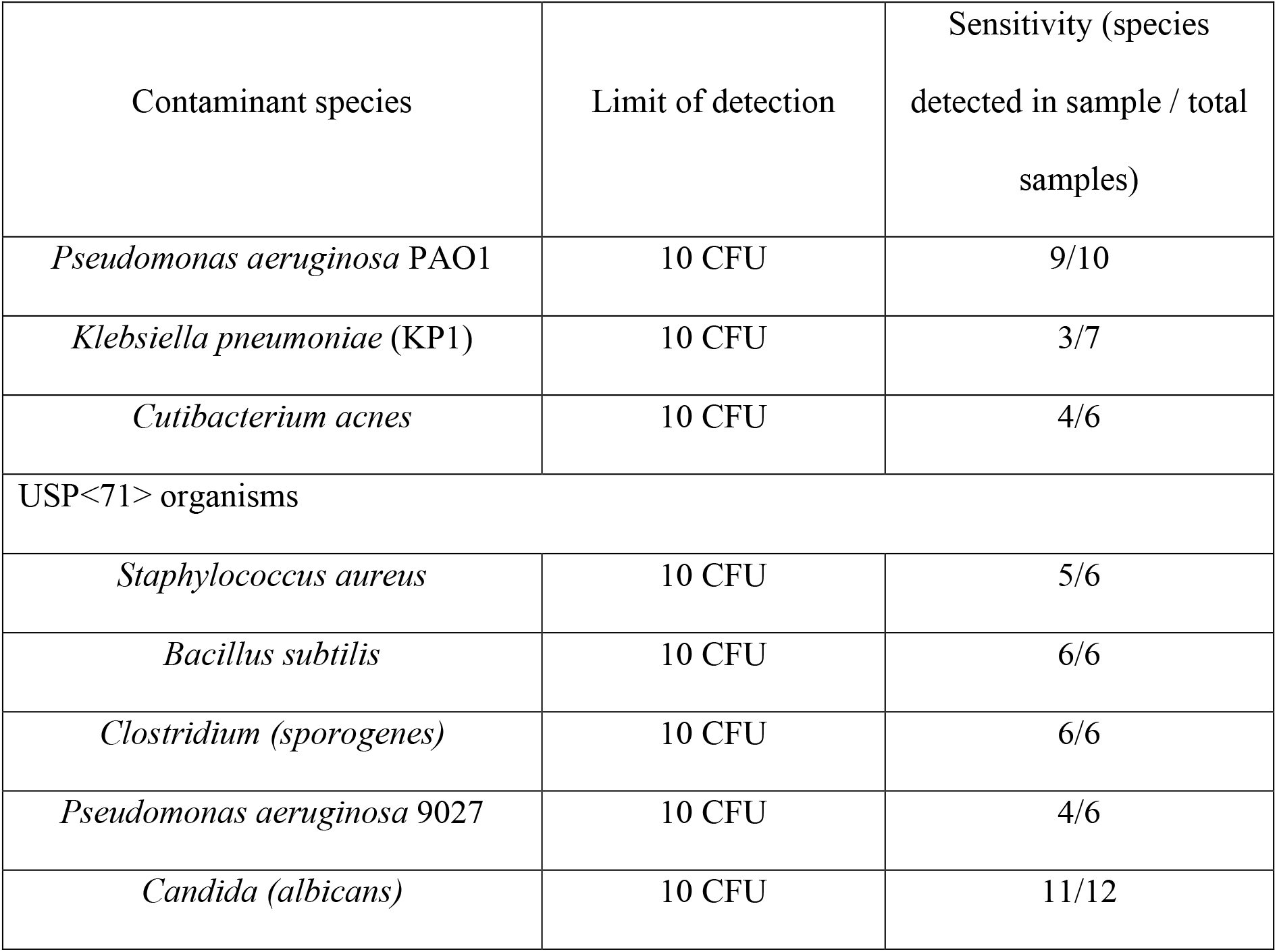
Limit of detection and sensitivity for contaminant detection. In the sensitivity column, the first value denotes total samples from amplicon sequencing split for combined spiked T- cell contaminant and microbial culture of that contaminant. Second value shows only the T- cell contaminant amplicon sequencing value.

| Contaminant species | Limit of detection | Sensitivity (species<br>detected in sample / total<br>samples) |
| --- | --- | --- |
| *Pseudomonas aeruginosa* PAO1 | 10 CFU | 9/10 |
| *Klebsiella pneumoniae* (KP1) | 10 CFU | 3/7 |
| *Cutibacterium acnes* | 10 CFU | 4/6 |
| USP<71> organisms |  |  |
| *Staphylococcus aureus* | 10 CFU | 5/6 |
| *Bacillus subtilis* | 10 CFU | 6/6 |
| *Clostridium (sporogenes)* | 10 CFU | 6/6 |
| *Pseudomonas aeruginosa* 9027 | 10 CFU | 4/6 |
| *Candida (albicans)* | 10 CFU | 11/12 |

The performance of both standard compendial tests and BacT/ALERT system can be further improved upon. A thorough analysis by England and colleagues reported an average 40 h time to detection for the BacT/ALERT compared to 53 h for compendial testing. Within the acceptable runtime (<144 h), 100/118 (84.7%) tested isolates were detected by the compendial USP <71> methods. When running the sterility tests using the BacT/ALERT alongside supplemental Sabourand dextrose agar (SDA) plates were incubated up to 360 h (15 d), 100% of fungi were detected, while USP<71> detected 95.8% of contamination events. The authors report that the majority of fungal isolates were detected within 144 h for the manual USP<71> methods, however with automated systems fungal detection could take longer to resolve and were only detected by manual inspection after 360 h of incubation. Additionally, the authors report that some bacterial species could not be detected without the infusion of fresh human blood into the culture e.g. *Haemophilus* influenzae and *Cutibacterium acnes* using the BacT/ALERT after 270 h. Furthermore, isolates grown at low inocula <30 CFU proved difficult to detect within the accepted time frame (<96 h, bacteria; <144 h, fungi). ^4^ In summary, within the accepted time frame for bacteria (<96 h), 83.1% of isolates were detected with the USP<71> compendial approach and 87% for the 32.5°C BacT/ALERT. Meanwhile for fungal isolates (<144 h), 87.8% of isolates were detected with the USP<71> compendial approach and 63.4% for the 32.5°C BacT/ALERT^4^. Given the described time constraints, as well as reduced sensitivity to filamentous fungal species, we propose an alternative methodology to detect and identify microbial contaminants through long-read MinION sequencing.

In the case of autologous CAR-T therapy products, each manufacturing lot is prepared for a single patient. Each lot of a cell therapy must pass through manufacturing release testing. This poses a limitation with regards to available test material – using a minimal volume for sterility assessment is important to maximise cells available for the patient, (with guidelines described in USP<1071>),^9^ while minimising additional manufacturing costs. The products will have in-process and final product testing for sterility, endotoxin levels, mycoplasma and replication competent virus.^10^ As CAR-T treatments become more readily available, rapid throughput and multiplexing will become increasingly necessary for the analysis of large numbers of scaled-out autologous cell therapy samples.

We propose an approach that identifies contaminants by the presence of either bacterial or fungal DNA. Long read MinION sequencing offers a rapid, simple-setup, real-time reads (contaminant identification before sequencing completion) and low-cost approach to DNA sequencing.^11, 12^ Compared to Illumina platforms, which can take days to weeks to complete sequencing and bioinformatics analyses, the Nanopore sequencing can provide results in less than 24 h.^13, 14^ Another advantage with the long read MinION sequencer, is greater taxonomic resolution than amplicon sequencing using the Illumina MiSeq system at the species level.^15^ This positions an amplicon sequencing approach using the long read MinION combined with a full size metagenome database favourably for managing rapid release testing.

Samples processed for T-cell therapy release tests will present at low sample volume and low contaminant DNA concentration. The long MinION offers long read sequencing with two potential routes for identification of contaminants: direct DNA and amplicon-sequencing. Direct DNA sequencing comes with the constraint of high background host DNA that will be sequenced; this can make detection of the relatively lower concentration microbial DNA harder. Autologous cell therapies will present as low volume low contaminant concentration samples, thus by necessity we took the PCR-based amplicon approach, which has the additional benefit of reducing microbe to host noise before downstream computational screening for host reads. In the case of bacteria, the highly conserved prokaryote 16S ribosomal RNA region is widely used in the metagenomics field with either the amplicon or shotgun metagenomics approach to perform microbial identification at the species level,^16–18^ while the equally highly conserved eukaryote 18S – 28S operon is used for fungal classification.

## Results

### Design of the DNA read analysis pipeline

The overview of the pipeline is described in Figure 1A, which processes reads derived from the spiked samples and derives a decision as to whether the sample is contaminated or not. The methodology for data acquisition, model training and validation, followed by the decision matrix are described in Figure 1B. In this analytical sterility study, we detected low levels of microbial contamination (10 CFU / mL) with high specificity and accuracy. The workflow we developed is compatible with low sample volume and rapid turnaround that meets patient needs and preserves sample shelf-life.

**Figure 1:**
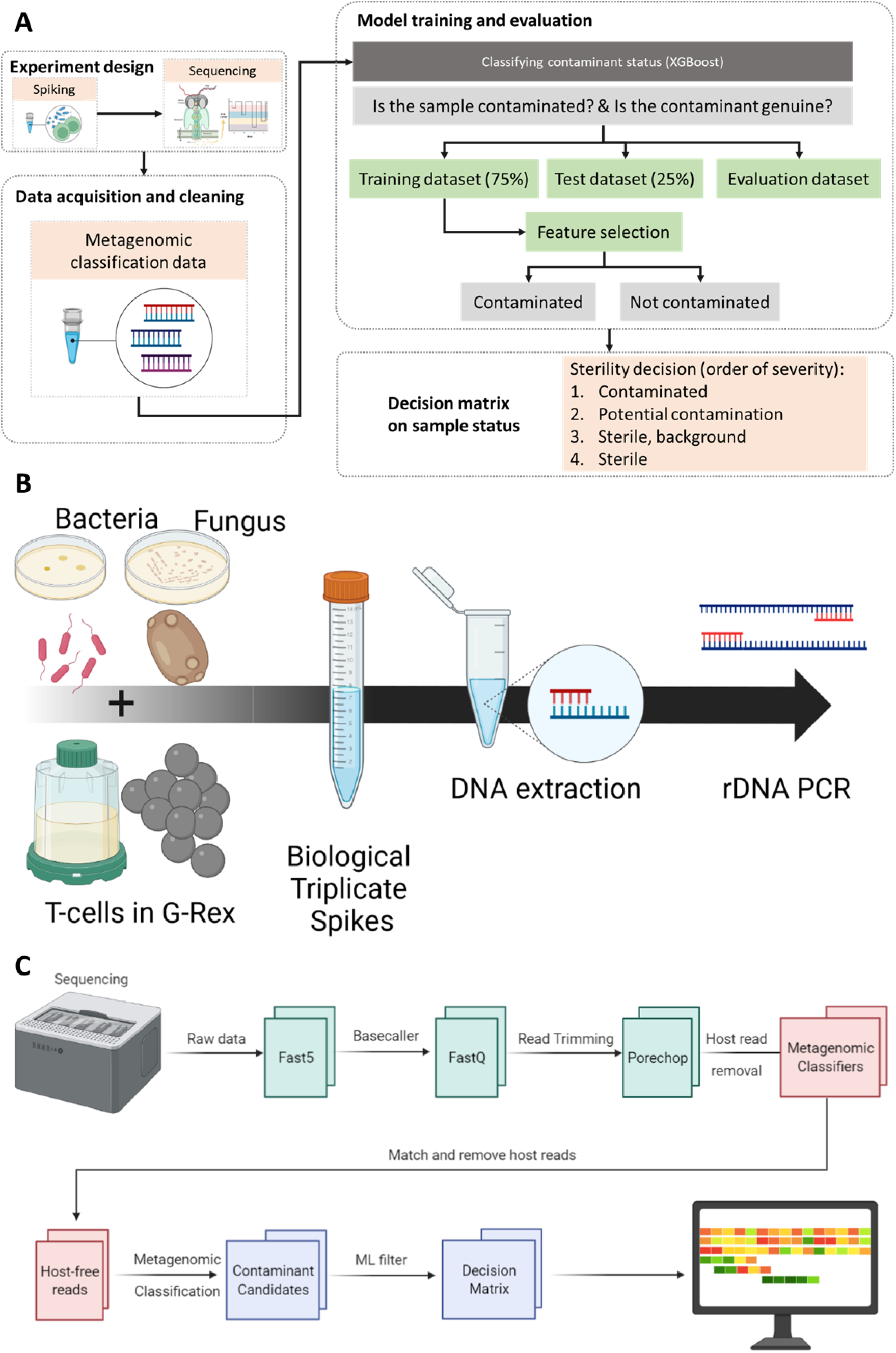
Pipeline workflow. **A.** Machine learning pipeline overview: microbial contaminants were prepared, sequenced and the reads processed. The metagenomic classification data, overall run read quality data, predicted species quality data and time to next read data were combined into a single table of features. A decision tree gradient boosting classifier algorithm, XGBoost was deployed to assess contaminant sterility status, for more information see Machine learning pipeline under methods or <u>github</u>. **B.** Bacteria (gram positive or gram negative) or fungus (yeast) are spiked into PBS-washed cultured T-cells. The process is repeated 3-fold with and without T-cells using cells from a different passage and separately cultured microbes. DNA is extracted using mechanical lysis, buffers and magnetic beads. DNA is amplified using targeted rDNA primers for the 16S region and 18S-28S region. **C.** Sequencing analysis pipeline: sequenced base called reads were cleaned and host reads removed. Remaining reads were classified against the combined viral, fungal and bacterial database using *Centrifuge* and *High-Speed BLAST*. Classified reads along with other data were provided to the machine learning pipeline for sample contamination status analysis.

Samples were comprised of negative samples (T-cell only, medium-only and cell-free medium) and target organisms (fungal and bacterial species) (Table 1). The table contains information about the added species, barcode used, sequencing time duration and the Nanopore kit used. Negative samples were prepared alongside inoculated samples, which were run in both direct and amplicon-sequencing runs. Only the amplicon-sequencing data is presented here.

### Inclusion of low-quality reads improved correctly classified read count

Our goal was to detect microbes at low concentrations (≤ 10 CFU). Detection therefore required generation of additional genetic information or a means of improving the pool of reads for assessment. We did not screen by phred quality score (Q) as we sought to identify contaminant read species identity. Consequently, there were a median of 6.9% additional reads across all samples. Mean read length was compared between high quality (Q ≥ 7) and high quality plus low-quality reads (Q > 0) for correctly classified true positive reads (Figure 2A). For 16S amplicon enriched species, the read length did not change greatly between the two groups, at around 1450 bp, while 18S – 28S had smaller mean fragment size (Figure 2A).

**Figure 2:**
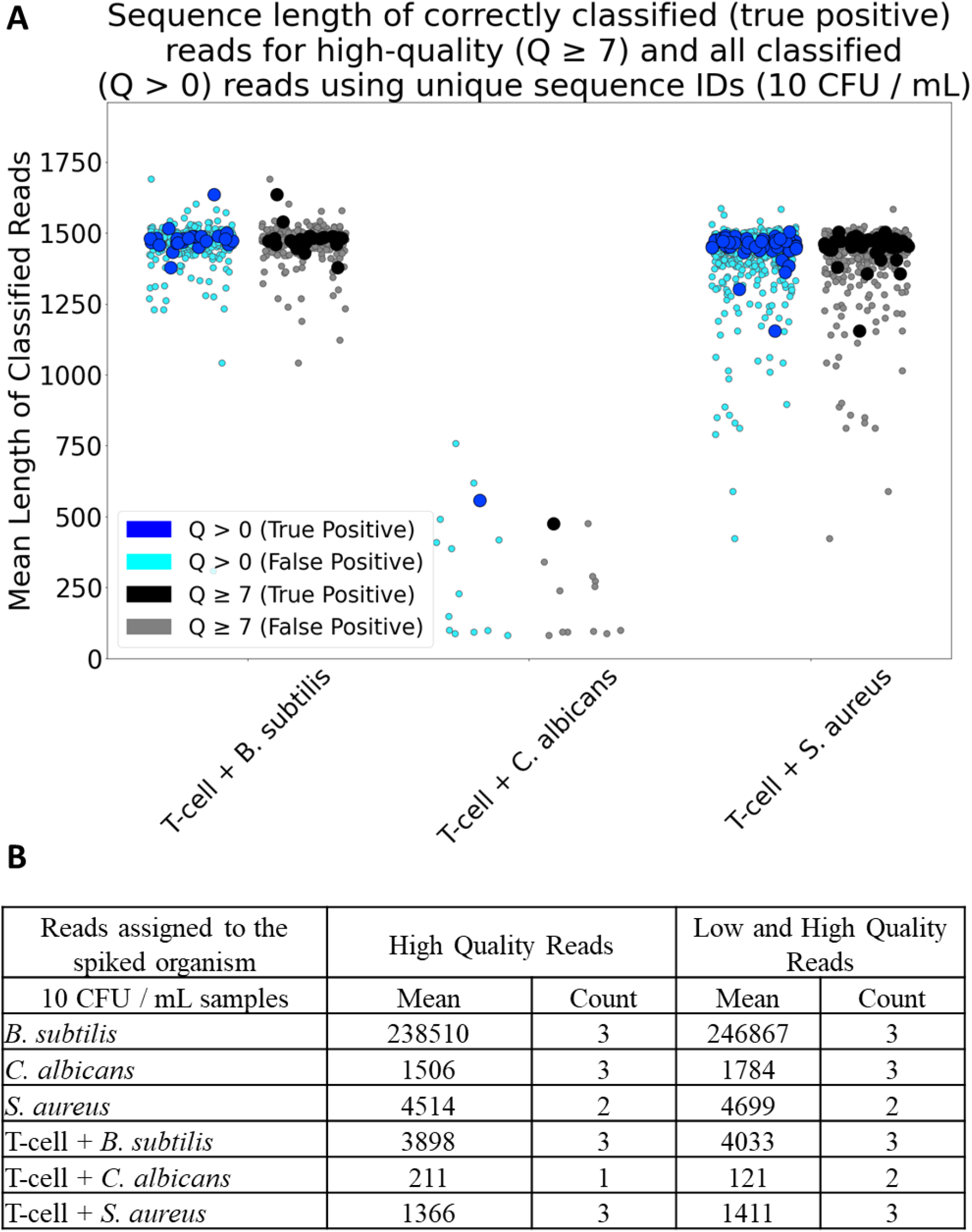
Assessment of incorporating low-quality reads in sequence classification. **A.** Additional low-quality reads incorporated into data analysis (N=3) for 10 CFU / mL samples. Mean read length for correctly classified (true positives) reads with and without use of the low-quality reads were depicted in blue and black, respectively. Incorrectly assessed reads (misclassified) were depicted in cyan and grey. **B.** Summary table of high-quality reality reads compared to using all reads for the microbes alone and microbes spiked into T-cells at 10 CFU / mL. Read numbers are for reads assigned to the correct spiked organism and represent a subset of all sequenced samples.

We calculated summary statistics for a subset of low concentration (10-100 CFU / mL) contaminated samples compared to microbe-only samples. Generally, we observed more microbe reads in the microbe-only cultures compared to samples containing host reads too; this was independent of low-quality read inclusion (Figure 2B). For example, by incorporating the low-quality reads, *Candida albicans* was detectable in two of the three samples contaminated at 10 CFU, while when using high-quality reads alone, *C. albicans* was detected in only one of the analysed samples. A side-by-side comparison of high-quality vs. any-quality reads revealed that by including the low-quality reads, one additional true positive sample was correctly identified; the overall read count increased for correctly predicted species. However, there was a concomitant increase in the number of misclassified species. As such, it is important to consider the use case before making the choice to include this additional source of reads.

### Detection sensitivity and time to detection

Initially, we sought to understand the limit of detection for a single species; *Pseudomonas aeruginosa* PAO1. The rationale for selecting a limit of detection of 10 CFU / mL is based on the USP<1071> chapter discussing infectious dose. The document discusses that a contaminant detection of 100 CFU / mL would catch all viable organisms, as such we set 100 CFU / mL as our target limit of detection, with the aim of detecting at lower concentrations as a form of stringent validation.^9^ T-cells were spiked with serial ten-fold dilutions of *P. aeruginosa* to the lowest input of 10 and 100 CFU / mL. Our approach examined the utility for both direct shotgun sequencing and amplicon sequencing. Extracted DNA was processed for both direct and amplicon sequencing. We consistently achieved 10 CFU / mL detection (from 1 mL spike sample) with the amplicon approach, while the limit of detection for direct sequencing was 1000 CFU / mL (data not shown). We then proceeded to contaminant detection for microbial cultures and T-cells for intentional contamination with *C. acnes, Klebsiella pneumoniae, Escherichia coli, P. aeruginosa*, or the USP<71> species *C. albicans, Staphylococcus aureus subsp. Aureus, P. aeruginosa, Bacillus subtilis subsp. Spizizenzii,* and *Clostridium sporogenes* (Figure 3A-B). In all cases, we consistently detected the contaminant to 100 CFU / mL (Figure 3A) and 10 CFU / mL (Figure 3B) in microbially contaminated T-cell cultures. One organism that proved consistently difficult to amplify was *K. pneumoniae*, resulting in fewer reads per sample than other species. Others have reported difficulty with taxonomic resolution in *Klebsiella* when amplifying the 16S rRNA regions of V1V2, V3 and V6V7 with Nanopore. They observed misclassification to closely related genera and low count of total mapped reads.^13^

**Figure 3:**
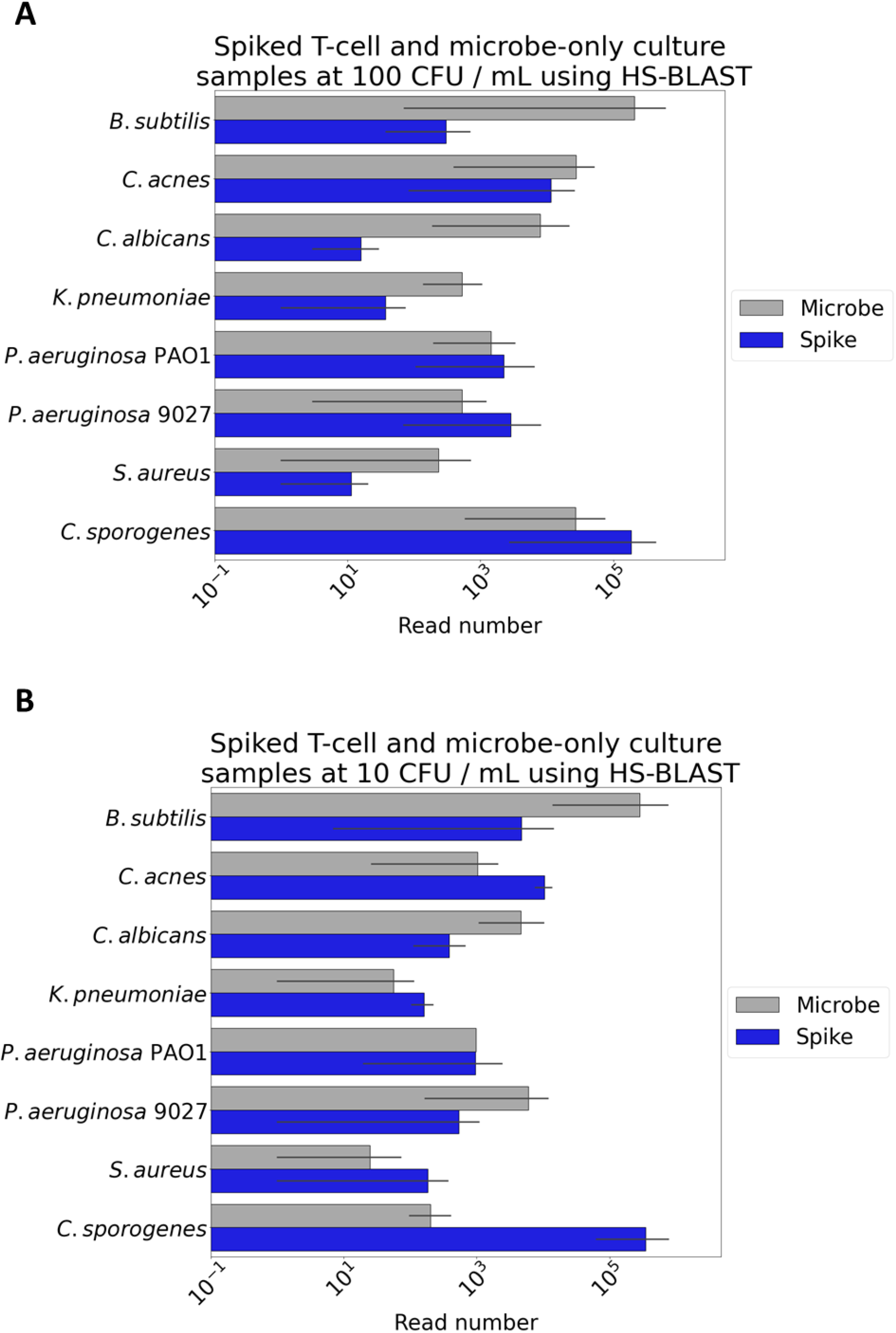
Microbes spiked into T-cell cultures. All samples were amplified with either 16S primers for bacterial species, or 18S-28S primers for fungal species. Samples were prepared for analysis as microbial cultures as well as simulated microbial contamination by the addition of microbes to activated T-cells at 100 CFU / mL **(A)** and 10 CFU / mL **(B),** pure culture spikes (grey) compared to contaminants spiked into T-cells (blue). Species tested were *K. pneumoniae, P. aeruginosa* PAO1, *C. acnes* and the USP<71> species; *C. albicans*, *B. subtilis*, *Clostridium sporogenes*, *S. aureus*, *P. aeruginosa* 9027. Error bars are biological replicates.

Our primary goal was to identify sample contamination as rapidly and accurately as possible. We were able to detect and identify contaminants within 24 h; including DNA extraction (2 h), PCR (2 h), 12 h for sequencing and the bioinformatics analysis (1 h). If the sequencing was shortened to 2 h, the time to detection was approximately 8 h per sample. However, this might not provide enough time to sequence enough reads dependent on the species present (in species that amplify well, e.g., *B. subtilis,* we observed enough reads within 2 h). Multiple biological samples can be processed simultaneously with multiplexed barcoding, allowing up to 12 samples to run on the same flow cell. This can be replicated five-fold for technical replicates per USP<1223> guidance,^19^ which can be run in parallel with the primer and culture medium negative controls. To further improve the time to detection, we evaluated different computational processing approaches. The read trimming and metagenomic classification were very time intensive. However, multiprocessing resulted in an average 4-fold improvement in speed. A computer system with higher specifications than used here would see an even greater improvement in time to result with more simultaneous processes.

### Machine learning pipeline and sample sterility status

The methods described in the pipeline allow for rapid and sensitive detection of low concentrations of contaminations when the contaminant is known. In addition to the proposed machine learning approach, all experiments would have plates cultured alongside the sequencing analysis. This would allow organisms that are detected as borderline to be captured by a secondary, albeit slower methodology. To detect unknown contaminants using an unbiased approach, we used machine learning to enable decision making regarding the sterility status of a sample. The metagenomic classifiers yield multiple potential species identities for perceived contaminants, which we initially attempted to deplete by using filters based on total minimum read count per species. Two separate pairs of models were generated for both data from Centrifuge and HS-BLASTn. The Centrifuge models assessing sterility status and whether a contaminant was correctly predicted are shown in Figure 4A-C. Model analysis for the HS-BLASTn model is depicted on Figure 4D-F. The classification report demonstrates a model that can identify sample contaminant status, while the model examining contaminant identity can find contaminant species (Figure 4A-B, 4D-E). The model predictions from sample status (sterile: true negative or contaminated: true positive) and correctness of contaminant classification (correctly classified vs misclassified contaminant) were combined and a decision matrix was used to decide if a sample was sterile, contaminated, potentially contaminated or showed signs of most likely being sterile with background noise signal (Figures 4C, 4E). For Centrifuge, 65.0% of negative controls were correctly labelled as sterile, 64.2% of 10 CFU / mL samples were correctly assessed as contaminated and 77.3% of 100 / mL CFU samples were identified as contaminated. Overall, while similar, the BLAST models perform better than Centrifuge on the final assessment for sterility status, with reduced ambiguity in a final decision on whether a negative control sample is sterile (75.0%) or whether a spiked sample was contaminated (10 CFU / mL; 83.9%, 100 CFU / mL; 96.0%). Overall, the 95% confidence intervals for the combined Centrifuge models were 80.67% ± 1.06% and 76.75% ± 1.03% for HS-BLASTn.

**Figure 4:**
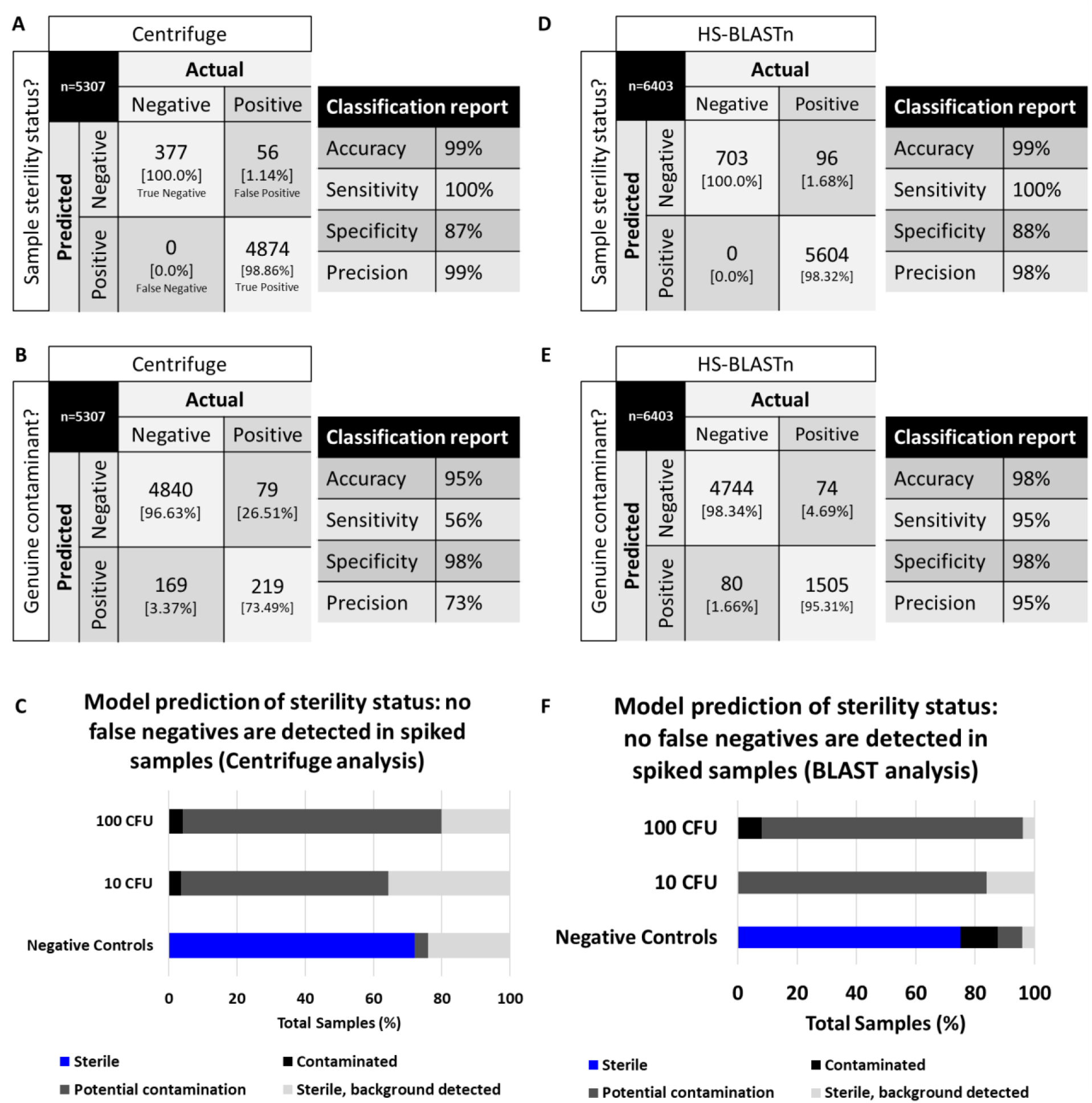
Machine learning XGBoost model performance, sample and prediction contaminant status. **A-C.** Model performance statistics from data from centrifuge metagenomic classifier used to generate two XGBoost classification models. **D-F**. Model performance statistics from data from high speed BLASTn metagenomic classifier used to generate two XGBoost classification models. **A, D.** Confusion matrix and classification report for model assessing sample sterility status. **B, E.** Confusion matrix and classification report for model assessing whether a predicted contaminant is a correctly classified contaminant or misclassified. **C, F.** All spikes and negative control model predictions were assessed for prediction accuracy regarding whether the sample assayed is sterile. Black bars depict samples assigned as likely contaminated, blue bars depict samples identified as sterile, while grey depicts samples where the algorithm had difficulty assigning a decision of either sterile or contaminated. CFU: colony forming units. Sample status is defined as sterile: true negative or contaminated: true positive. Correct contaminant classification is defined as a true positive contaminant vs a misclassified contaminant.

## Discussion

We have demonstrated that amplicon sequencing can be used for low sample volume (< 1 mL), low concentration (≤ 10 CFU) microbe detection in intentionally contaminated T-cell cultures. Our goal was to detect a contaminant with high specificity and in a short turnaround time to enable rapid sterility release testing for cell therapy products. We designed our sterility detection to be faster than current commercial detection times for low concentration contamination events of < 100 CFU / mL to within < 24 h, which compares with the historical detection times of 7 – 14 d using BacT/ALERT^20, 21^ and more recent detection times of 40 h,^4, 9^ which is in line with FDA sterility guidelines.^22, 23^ For bacterial and fungal detection, we used an amplicon approach that significantly increased the contaminant signal from DNA extraction. Use of the Nanopore 16S amplicon kit allowed us to generate consistent full length 16S fragments. Previously, full length amplicons have been shown to be comparable in reliably identifying genera on Nanopore and MiSeq technologies, while Nanopore can operate at lower run costs (50 USD / sample).^14^ However, when generating similar fragments using the 18S-28S primers, we did not observe similar read length consistency. This is likely due to use of the transposase technology in the Rapid Barcoding Sequencing Kit (SQK-RBK004) that cleaves the PCR product for barcode ligation. The problem could be solved by using the Nanopore ligation kit (SQK-LSK110), however this approach increased sample preparation time by up to 5 h. This limited our ability to sequence the entire region as a single strand, however as we have demonstrated, we are still able to detect *C. albicans* to 10 CFU.

During the sequencing analysis, we observed background genomic material in the sterile media controls. We believe these are derived from DNA contaminants in the DNA extraction, PCR and library preparation kits. The presence of a kitome has been widely described in the literature, including from the DNeasy PowerSoil Kit used in this work.^24, 25^ This is problematic because at low microbial concentration the signal to noise ratio becomes elevated, making identifying correctly classified contaminants difficult and increasing read misclassification. False positive predictions will potentially disrupt the ability to deliver the cell therapy product to a patient in the needed timeframe because of additional time required to carry out failure investigation to assess actual lot sterility status, in addition to incurring further manufacturing costs. The issue of high noise at low concentration reinforces the need to use negative controls to identify the contamination background as well as control the number of PCR cycles, as previous studies have shown that 20 cycles were too few while 40 cycles will amplify the noise.^26^

One of the key limitations of this approach is the use of DNA for detection of adventitious agents. This does not provide a definitive answer to the viability of the detected contaminant and is part of the reason a deeper understanding of the kitome is required. Alternatively, sequencing of RNA after reverse transcription and rDNA amplification would provide a means to test for contaminant contamination directly. We are investigating background species that are detected during the sequencing, they are characterised by weak signal and may be on the limit of perception due to the low read count derived from an already very low starting sample concentration and volume. Extensive testing and validation of potential background contaminant species could reduce noise detection, we propose performing multiple technical replicates per sample, as well as running multiplexed positive and negative standard controls in every sterility test.

To optimise sensitivity and pipeline runtime, we examined the choice of input species to build the metagenomic databases. This is mostly an issue with the NCBI fungal refseq, which at the time of database construction was confined to only 12 complete genomes. To expand the range of species, we incorporated incomplete genomes for both the *BLASTn* and *Centrifuge* databases. However, the bacterial reference database is very large and contains many genome replicates for common bacterial genera *Pseudomonas* (613) and *Escherichia* (1100), as well as species including *P. aeruginosa* (219) and *E. coli* (1078). As such, to reduce the database size, we retained only genomes with a “complete genome” and removed those labelled as “chromosome” or “plasmid”. This helped improve runtime and reduce database size differences between the fungal and bacterial databases. The only downside we observed by shrinking the bacterial database was the loss of species-level inference as observed with *C. sporogenes*, where *C. butyricum* was indicated instead. For future organisms, a living database could be constructed and periodically updated. Reference genome agnostic approaches could be used for unknown contaminants, such as read binning using the DNABERT approach described below.^27^ After inclusion of *C. sporogenes*, we continued to observe that *C. butyricum* was preferentially predicted with 42% more reads on average for *HS-BLASTn* (and not identified by *Centrifuge*).

We subtracted host sequences using the human genome to reduce noise in the analysis pipeline and improve pipeline efficacy. With this approach, 98% of total reads were identified as host reads and removed, illustrating the challenge of detecting low level bacterial or fungal contaminants in samples of cell culture material. Our study focused on amplicon-sequencing, which enriched for target sequences and reduced the ratio of host reads to target reads. We implemented host read removal to improve sensitivity and enable a non-amplicon approach going forward. Nevertheless, host read removal is useful for improving microbial identification, as has been described previously in viral contaminant detection.^28^ Previously, host read removal has been shown to improve viral contig assembly while reducing the number of assembled contigs so as to allow use of metagenomic classifiers such as *BLASTn*^29^. In addition to removing host reads, removal of background reads by sequencing healthy and sterile T-cells, then subtracting the reads from the final pool could further improve sensitivity by reducing misclassification events.^29^ The 16S and 18S-28S amplicon sequencing approach is more successful than direct metagenome sequencing, especially at contaminant concentrations ≤ 1000 CFU / mL. Previous studies have demonstrated that amplicon-based approaches can introduce inaccuracy and misclassification, for example 0.93% of reads are misclassified by EPI2ME at the genus level^16^ and that there is a 2.09% misclassification for the NCBI 16S reference database at genus level.^16^ For cell therapy sterility, the improved contaminant detection at the cost of the a small increase to misclassification is an acceptable outcome.

Two key limitations for model development are the volume of available sequencing analysis data, as well as the high sample-to-sample variance in the data distribution. For example, we observed sources of variation from read count, read quality and experimental runtime. The implications for the observed variance include difficulty in applying the machine learning model to new samples that may contain data with novel distributions. Solving the problem of noisy, unreliable data requires a robust and flexible model that can classify despite lack of knowledge on the data distribution for the evaluation samples. This includes selecting for features with minimal missing data and applying, as well as deploying data augmentation techniques such as addition of gaussian noise, generating squares and taking log10 of the numerical data. Finally, we have applied regularisation techniques during model development including optimising L1 (lasso regression), L2 (ridge regression) and the gamma hyperparameters to mitigate overfitting. To improve issues with generating more similar and reliable samples that follow a specific data distribution would require further refinement of the DNA extraction and library prep, in addition to controlling for missing data points during feature selection. However, this would come at the cost of having a model that is less able to generalise to other cell types.

Our analysis has primarily focused on generating many instances of contaminated samples. During production for cell therapy manufacturing, we envisage sample contamination events as rare. Consequently, future model optimisation must be considered through this data imbalance, as observed in fraud detection for better anomaly prediction.^30–32^ Alternatively, it might be possible to replace the metagenome classifier and binary classifiers with a single step natural-language processing-based (NLP) approach using e.g. DNABERT.^27^ A NLP-based approach would complement the metagenomics analysis that we have already designed. Furthermore it would give us a potential solution for the identification of unknown contaminated samples independent of whether they have been previously characterised.

In conclusion, we have developed a rapid (< 24 h), reliable, sensitive and specific long-read sequencing pipeline for the detection of microbes in T-cell therapies alongside a large dataset of microbial organisms (USP <71> focus) low concentration samples in pure culture and spiked with T-cells. This is partly achieved by using machine learning to provide an unbiased untargeted approach that permitted automated decision making of the sterility status of cell therapy products. Our approach has demonstrated that we can achieve high sensitivity and detect contaminated samples on par or superior to accepted methods in superior time frames.

## Materials and Methods

### Cell culture

Healthy human primary T-cells (Human PBMCs were obtained from commercial leukopak (Cat#260240.01, Lonza)) were cultured in AIMV (Cat#12055091, Gibco)+ 2% AB Human Serum (Cat#H4522, Sigma-Aldrich) + 100U/mL IL-2 (Cat#130-097-743, Miltenyi Biotech) and activated using ImmunoCult™ Human CD3/CD28 T-Cell Activator (Cat#10971, StemCell Technologies). T-cells were cultured for 14-21 d. Cell counting was performed using a TC20™ Automated Cell Counter (Biorad) with Trypan blue stain.

Microbial species were grown and cultured as described in Table 1. We used and developed standard curves to accurately estimate the number of bacterial cells present within a culture at a given timepoint using the optical density (OD) values.^33^ Colony forming units (CFUs) were counted before the addition of microbial cells to sterile cultured T-cells. Briefly, 100 μL culture was serially diluted, 5 μL of each serial dilution was plated on agar, and CFUs were counted the following day. For low concentration samples, following overnight culture, a subculture was prepared at 0.05 OD and incubated for 1-3 h (the time was determined using the standard curve to predict the log phase of growth) and cultured to 0.5 OD. The culture was then diluted to 0.1 OD and subsequently serially diluted with LB and plated on agar plates for CFU counting.

### Preparation of spiking samples

Spiked samples were prepared using activated T-cells and bacterial or fungal species. Each sample used 500,000 T-cells. Bacterial and fungal cultures at a range of 10-100 CFU / mL were prepared as described above and added directly into T-cells in PBS and incubated for 5 min.

### DNA extraction and amplification

For sample preparation, a uniform approach to DNA extraction was taken to maximise DNA extraction from all possible microbes. Using the manufacturer’s protocols, the DNeasy PowerSoil Pro Kit (Cat#47014, Qiagen) was the most consistent and reliable product. The kit disrupts cell walls and membranes alike through bead beating (TissueLyser II [Cat#85300, Qiagen]). The samples were quality controlled using the NanoDrop™ 2000/2000c Spectrophotometers (Cat#ND-2000, Thermofisher Scientific) to assess RNA and protein contaminants (260/280 (1.8-2.0), 260/230 (2.0-2.2)) and Qubit 4 Fluorometer (Cat#Q33238, Thermofisher Scientific) for DNA concentration.

Primers to amplify full length18S-28S rRNA genes were adapted from^34^ to amplify the entire operon. 18S NS1 short F: CAGTAGTCATATGCTTGTC and 28S RCA95m R: CTATGTTTTAATTAGACAGTCAG.^34^ All 16S primers described were those available in the Oxford Nanopore 16S-barcoding kit (SQK-RAB204). PCR conditions were as follows: initial denaturation 1 min at 95°C for 1 cycle, 20 s denaturation at 95°C, for 29 cycles, 30 s annealing at 55°C for 29 cycles, an extension time of 2 min at 65°C for 29 cycles, followed by a final 5 min extension at 65°C. LongAmp Taq 2X Master Mix (e.g. NEB M0287).

### Long read sequencing library preparation

We used a multiplexing approach to maximise the sample throughput. This uses the existing 16S primers that have barcodes associated with them that can later demultiplexed. In the case of the 18S-28S amplicons, no pre-existing kit exists. Thus, we applied barcodes from the rapid barcoding kit (SQK-RBK004) to the 18S-28S amplified samples, which could later be demultiplexed. Briefly, amplified DNA had RB01-12 fragmentation primers attached and were concentrated as per the manufacturer’s protocol. The Oxford Nanopore MinION sequencer with a MinIT device for base calling. The FLO-MIN106 flow cell was used to run the DNA samples for between 2 – 24 h. Reads were generated and basecalled (the electrical impedance signal was converted to a kmer string) by the Nanopore device were processed using the basecaller *Guppy* v3.2.10.

### Pipeline tools

Metagenomic classification databases were generated from NCBI bacteria, virus and fungal databases, these were combined into a single database. Due to runtime issues with the bacterial database using BLAST, we developed an abridged database “*filter-bacteria*” that contained a reduced number of entries for highly prevalent organisms within the NCBI database, e.g. *E. coli*. This reduced the database to 1/6 its original size and was combined with the viral and fungal sequences to make a viral-fungal-bacterial database. To augment the fungal database for *HS-BLAST*, additional genomes were included from NCBI that were labelled as incomplete genomes. The database location and information were saved in a .txt file. Three metagenomic classifiers used in the pipeline were: *Centrifuge,*^35^ *high-speed (HS-) BLASTn*^36^ and *Krakenuniq.*^37^ The DNA reads from sequencing were used as the raw data for the metagenomic classifiers. Using “*centrifuge-download*”, the NCBI refseq libraries for viruses (24/06/2020), fungi (25/05/2021) and bacteria (21/09/2020) were acquired and “*centrifuge-build*” was applied to generate the database using the NCBI taxonomy structure. *Krakenuniq* was an additional classifier used for removing the maximum number of host reads, in order to prevent the host reads being a majority. *Porechop* 0.2.4 (https://github.com/rrwick/Porechop) trimmed adapters from the Nanopore reads. *Qcat* 1.1.0 demultiplexed the barcoded reads, as well as the updated guppy algorithm (v6.1.5).

### Pipeline

The bioinformatics pipeline (Figure 1A, B) links together multiple tools, in addition to hosting custom scripts to check analysis progress. If the sample was multiplexed, it was demultiplexed before read trimming by *porechop*. We retained low quality reads from the “*fastq_fail*” folder within the pipeline because the aim was to detect low concentration contaminants.

If the user specifies a known host species and reference genome, (e.g., human), the pipeline will align host reads using *HS-BLAST*, *Krakenuniq* and *Centrifuge*. The read IDs were used to remove these host reads from the read pool, which accelerates the workflow and reduces noise related to the host reads. Host read removal was completed in tandem using multiprocessing for each metagenomic classifier. Statistics on the number of host reads, classified and unclassified reads were retained. Host-depleted reads were then processed using the metagenomic classifiers against the combined fungal, viral and bacterial database. *HS- BLAST* and *Centrifuge* output were used to generate descriptive summary statistics from the troubleshooting files using the pandas describe function. Summary statistics for each predicted contaminant were generated from the Guppy file “*sequencing_summary.txt*” by aggregating the individual read quality scores that were processed using the describe function.

After analysis using the pipeline above, *Nanoplot*^38^ was used to generate and monitor run statistics. A wrapper was created to generate unique sequencing summary files for barcoded samples by splitting the reads on their barcode IDs. The “*NanoStats.txt”* file was cleaned up for later use in the machine learning section of the pipeline. Run environment specification: Python version: 3.8.3, Bash version: GNU bash, version 4.4.20(1)-release (x86_64-pc-linux-gnu).

### Machine learning pipeline

Initial pre-processing for metagenomic classification data from the combined viral, fungal and bacterial database uses an independent filter for each metagenomic classifier. The pre-processing step is necessary because it filters out many of the low quality predicted species made by both *HS-BLAST* and *Centrifuge*. The *HS-BLAST* filter uses a maximum percent identity value greater than 83%. This reduced misclassified species by 33.19% (incorrectly predicted species removed from downstream analysis), while retaining (166/167) of samples, resulting in the loss of one sample from the analysis. Similarly, the *Centrifuge* filter uses a minimum metagenomic classification mean score greater than 900. The filter removes misclassified species by 38.64%, while retaining 97.45% (153/157) of samples. The *Nanoplot* output included experiment level summary information were subsequently concatenated to the data. Summary statistics using the *describe()* function were generated for the metagenome classification data. Predicted species that have *NaN* values (Not a Number) for read count and mean read quality were removed from the dataset.

The data were pre-split based on two layers of *train_test_split()* 75:25 split ratio. The first split generated the training and testing dataset alongside an unseen and (for model assessment) unused evaluation dataset. The training and testing were then split again into training dataset and test dataset. Features were selected using *Featurewiz* (https://github.com/AutoViML/featurewiz), *Featurewiz* was used to generate a list of important features that accounted for highly correlated features, as well as less insightful features and provided features for the generation of a high-performing model.

Binary classifiers were designed to answer two questions: (1) is the sample contaminated? and (2) is the predicted contaminant correctly classified and does it match the spiked species? For (1), the labels and encoded labels were as follows: *True_positive*: 1, *True_negative*: 0 and (2) *False_positive*: 0, *True_positive*: 1. *XGBoost* classifier models were developed and implemented. The data preparation was similar for each. *Gridsearch* was used to identify the ideal model starting parameters. Performance was assessed with cross validation testing (*cv=5, scoring=accuracy*), confusion matrices and generation of a classification report. Models were described below:

1. Is the sample contaminated? Defined as either sterile or not sterile.
2. Is the predicted contaminant correctly classified? Defined as expected contaminant from the spike or any other predicted species.

The model predictions were then compared to the original labels and compared for accuracy. This step was completed for both questions (1) and (2). The last step for the training and testing of the machine learning pipeline were to take the predictions and make an assessment using a decision matrix as to whether a given sample is sterile or contaminated based on the available data.

To evaluate the machine learning model, the trained machine learning model applied to an unseen data pool that shared the same distribution as the complete dataset: the evaluation data. The standard scalers were reloaded from the training step and used for standardisation. The features identified by *Featurewiz* were also imported and reused when running the model with new data.

## Supporting information

Supplementary Table 1 Samples

Supplementary Table 2 Centrifuge metagenomic summary

Supplementary Table 3 HSBLASTn metagenomic summary

Supplementary Table 4. Sequencing Summary Statistics

Supplementary Table 5. Sequencing Summary Statistics following metagenomics analysis

Supplementary Table 6. Rationale for choice of XG Boost

## Acknowledgments

This research is supported by the National Research Foundation, Prime Minister’s Office, Singapore under its Campus for Research Excellence and Technological Enterprise (CREATE) programme, through Singapore MIT Alliance for Research and Technology (SMART): Critical Analytics for Manufacturing Personalised-Medicine (CAMP) Inter-Disciplinary Research Group. Financial support from National Research Foundation and Ministry of Education Singapore under its Research Centre of Excellence Program was provided through Singapore Centre for Environmental Life Sciences Engineering. Graphical abstract and parts of Figure 1A, 1B and 1C were created with BioRender.com.

## Author Contributions

JS: Conceptualisation, Methodology, Validation, Software, Writing, Visualisation, Formal Analysis. MN: Validation, Investigation. WXS: Investigation. EL: Methodology, Validation. PWB: Writing - Review & Editing. JMW: Writing - Review & Editing. RBHW: Methodology, Writing - Review & Editing, Supervision. SAR: Conceptualisation, Methodology, Writing - Review & Editing, Supervision. SLS: Conceptualisation, Methodology, Writing - Review & Editing, Supervision.

## Conflicts of Interest

The authors have declared no competing interests.

## Data Availability

Pipeline can be found at (currently a private GitHub): https://github.com/Electrocyte/adventitious-pipeline/

Reviewer link for metadata (Sequence Read Archive (SRA) ID: PRJNA869859): https://dataview.ncbi.nlm.nih.gov/object/PRJNA869859?reviewer=l69tlml6fv0vhje0mbq1j476u6

## Notes

### Summary of Updates

This version has been updated to include additional graphical information in Figure 1, specifically by further dividing the figure to allow description of the entire workflow. We have reduced repetition within the main text body too where relevant. We have included additional background information to the introduction to clarify existing compendial methodologies as compared to our approach. Author list updated to reflect the contributions of all authors to the work. No changes made to supplemental files.

